# CryoEM Structure of the human THIK-1 K2P K^+^ Channel Reveals a Lower ‘Y-gate’ Regulated by Lipids and Anaesthetics

**DOI:** 10.1101/2024.06.26.600475

**Authors:** Karin EJ Rödström, Bisher Eymsh, Peter Proks, Mehtab S Hayre, Christian Madry, Anna Rowland, Simon Newstead, Thomas Baukrowitz, Marcus Schewe, Stephen J Tucker

## Abstract

THIK-1 (*KCNK13*) is a halothane-inhibited and anionic lipid-activated Two-Pore Domain (K2P) K^+^ channel implicated in microglial activation and neuroinflammation, and a current target for the treatment of neurodegenerative disorders such as Alzheimer’s and Amyothropic Lateral Sclerosis (ALS). However, compared to other K2P channels, little is known about the structural and functional properties of THIK-1. Here we present a 3.16 Å resolution cryoEM structure of human THIK-1 that reveals several unique features, in particular, a tyrosine in M4 (Y273) which contributes to a lower ‘Y-gate’ that opens upon activation by physiologically-relevant signalling pathways. We further demonstrate that binding of linoleic acid within a modulatory pocket adjacent to the filter also activates THIK-1, and that halothane inhibition involves a binding site within the inner cavity resulting in changes to the Y-gate. Finally, the extracellular cap domain contains positively-charged residues that line the ion exit pathway and which contribute to the unique biophysical properties of this channel. Overall, our results provide important insights into the structural basis of THIK1 function and identify distinct regulatory sites that expand its potential as a drug target for the modulation of microglial function.

## Introduction

Two-Pore Domain K^+^ (K2P) channels are a distinct subset of K^+^ channels that assemble as dimers to form a pseudo-tetrameric central pore. There are 15 human K2P (*KCNK*) channels that respond to a range of physical, chemical and biological signals to influence the resting membrane potential and couple these inputs to changes in cellular electrical activity^1,2^. Their functional roles, especially within the central and peripheral nervous system, means they represent important pharmacological targets and their dysfunction results in a variety of different disease states and neurodevelopmental disorders^3^.

THIK-1 is encoded by *KCNK13,* but in contrast to its more widespread expression in rodents^4^, its expression in humans is restricted to microglia^5^, the brain’s innate immune cells. As part of their role in protecting the brain from injury and invading pathogens, microglia can become ‘activated’ to trigger an immunological response aimed at containing neuronal damage^6–8^. However, in some cases, microglia may also adopt disease-exacerbating states of activation leading to neuroinflammation which is a key driver of neurodegenerative disorders such as Alzheimer’s, Parkinson’s and ALS (aka Lou-Gehrig’s Disease). THIK-1 channels have been shown to be critical for the release of pro-inflammatory cytokines during the activation of human microglia and *KCNK13* is upregulated in both animal models of neurodegeneration and Alzheimer’s Disease itself^9^. Furthermore, THIK-1 inhibitors slow the progression of neurodegeneration in animal models^10^. Likewise, THIK-1 activators may also help promote immune surveillance of the brain parenchyma in situations where microglial immune surveillance is partly compromised. Consequently, this channel represents an important therapeutic target for the modulation of microglial function^11^ and Phase 1 clinical trials of a THIK-1 inhibitor are currently underway^10,12^.

In comparison to many other K2P channels, the structural and functional properties of THIK channels are poorly understood. This subfamily of K2P channels were first identified in 2000^4^ as the TWIK-related-Halothane-Inhibited K^+^ channels, THIK-1 and THIK-2; yet despite robust functional expression of THIK-1 in heterologous systems, relatively few studies have focussed on this channel over the past two decades, consequently the structural basis for its biophysical and functional properties remains largely unclear. The other member of this subfamily, THIK-2 shares 62% sequence identity with THIK-1 and can coassemble with THIK-1 to form novel heterodimeric channels^13^. However, THIK-2 is largely retained within the endoplasmic reticulum (ER) thereby limiting its functional analysis; measurement of THIK-2 currents have therefore only been possible by removal of this ER-retention motif to promote trafficking to the cell surface and/or the introduction of mutations within the transmembrane helices that have an activatory gain-of-function effect^14,15^.

Recent studies have demonstrated that THIK-1 can be transiently activated by both G_i/o_ and G_q_-coupled receptor pathways^16^, and can also be directly activated by anionic lipids such as PIP_2_ and oleoyl-CoA^17^. In addition, THIK-1 exhibits direct inhibition by the phosphodiesterase inhibitor, 3-isobutyl-1-methyl-xanthine (IBMX)^18^ as well as several small molecule inhibitors currently being developed as therapeutics^10,12^. However, the detailed mechanisms of this pharmacology are not well understood, neither are its biophysical properties because the single channel conductance of THIK-1 is very low (<5pS) with only very brief flickery openings (<0.5ms) thereby limiting detailed biophysical analysis of its gating^19^. Given the importance of THIK1 as a potential therapeutic target, a greater understanding of its structural and functional properties is therefore warranted.

Here we present a cryoEM structure of human THIK-1 at 3.16 Å resolution in combination with a detailed functional analysis. This study reveals several unique structural features that provide important insights into the molecular basis of THIK-1 function, its regulation by lipids and volatile anaesthetics, and its suitability as a drug target for the regulation of microglial function.

## Results and Discussion

### CryoEM structure of THIK-1 reveals unique structural features

Similar to previous approaches used for K2P channels, we first removed the predicted unstructured C-terminal region of THIK-1 and confirmed this truncated channel (Gly^9^-Gly^297^) was functionally active. The channel was expressed in insect cells and purified for single particle cryoEM. The resulting structure was resolved to 3.16 Å resolution (**Figure 1**, **Supplementary Table 1** and **Supplementary Figure S1**).

**Figure 1.**
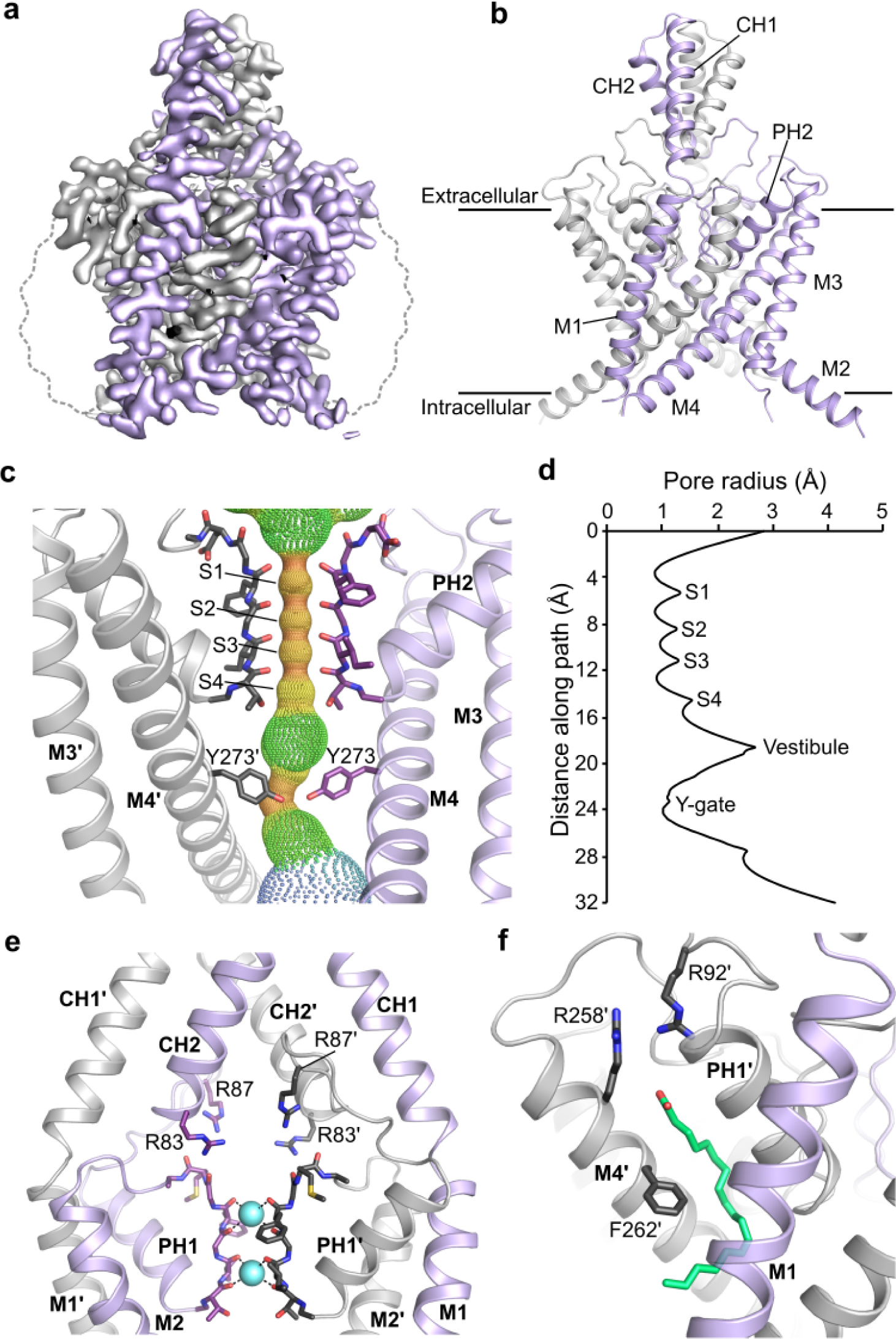
Structure of THIK-1. **a,** Sharpened cryoEM map viewed from the side with the density for THIK-1 channel subunits in gray and purple. The approximate position of the detergent micelle is outlined as a dotted gray line. **b,** Structure of THIK-1 coloured as in Panel a with the M1-M4, Pore and cap helices labelled. **c,** HOLE profile through the channel pore with the selectivity filter (S1-S4 sites) and the constriction formed by the lower tyrosine 273 (Y-gate) site highlighted. For clarity, M1, M2 and PH1 have been hidden. **d,** Pore radius of the THIK-1 channel interior as distance function along the ion permeation pathway. **e,** Ion exit pathway at the extracellular site of the selectivity filter is lined by a cluster of positively-charged residues (R83 and R87 from both subunits). M3, M4 and PH2 have been hidden for clarity. **f,** The K2P modulator pocket, showing a lipid bound at the inter-subunit interface of M4, PH1 and M1. Key residues in close proximity are highlighted as sticks.

The overall fold of the channel is similar to all other known K2P channel structures; it forms a domain-swapped homodimer, with each subunit containing four transmembrane helices (M1–M4), two pore helices (PH1 and PH2), two selectivity filter (SF) motifs (SF1 and SF2) and two extracellular cap-forming helices (CH1 and CH2) though it lacks a cysteine at the relevant position at the apex, so unlike many other K2Ps there is no interchain disulphide at this point. The SF adopts a near four-fold symmetry and is similar to most other K2P channels, but the intracellular M2-M3 loop (Arg^166^–Gly^189^) is longer than in most other K2P channels and was not resolved, presumably due to its conformational heterogeneity. (**Figure 1a-b**).

However, despite these similarities, several unpredicted structural features were identified which help explain the unique functional properties of this channel. Firstly, a lower constriction of the permeation pathway within the inner cavity created by the M2 and M4 pore-lining helices (**Figure 1 c-d**), a cap-domain with positively charged residues lining the extracellular ion exit pathways (**Figure 1e**), and a single chain lipid bound within the cryptic ‘K2P modulator pocket’ adjacent to the SF (**Figure 1f**).

### A lower ‘Y-gate’

Several K2P channels gate exclusively within the SF and do not possess a lower cytoplasmic gate. However, this structure of THIK-1 reveals a major constriction within the inner cavity is created by the interaction of bulky side chains on both the M2 and M4 helices; in particular a tyrosine (Y273) on each M4 helix points inwards to occlude the pore along with I139 on M2 (**Figure 1c and Figure 2a**). This constriction differs to the lower ‘X-gate’ found in TASK channels which only involves residues on M4 and is positioned slightly higher than the X-gate creating an even smaller inner vestibule below the SF. This constriction completely occludes the permeation pathway in THIK-1 (**Figure 1d**) and so this conformation is predicted to be closed and non-conductive.

**Figure 2.**
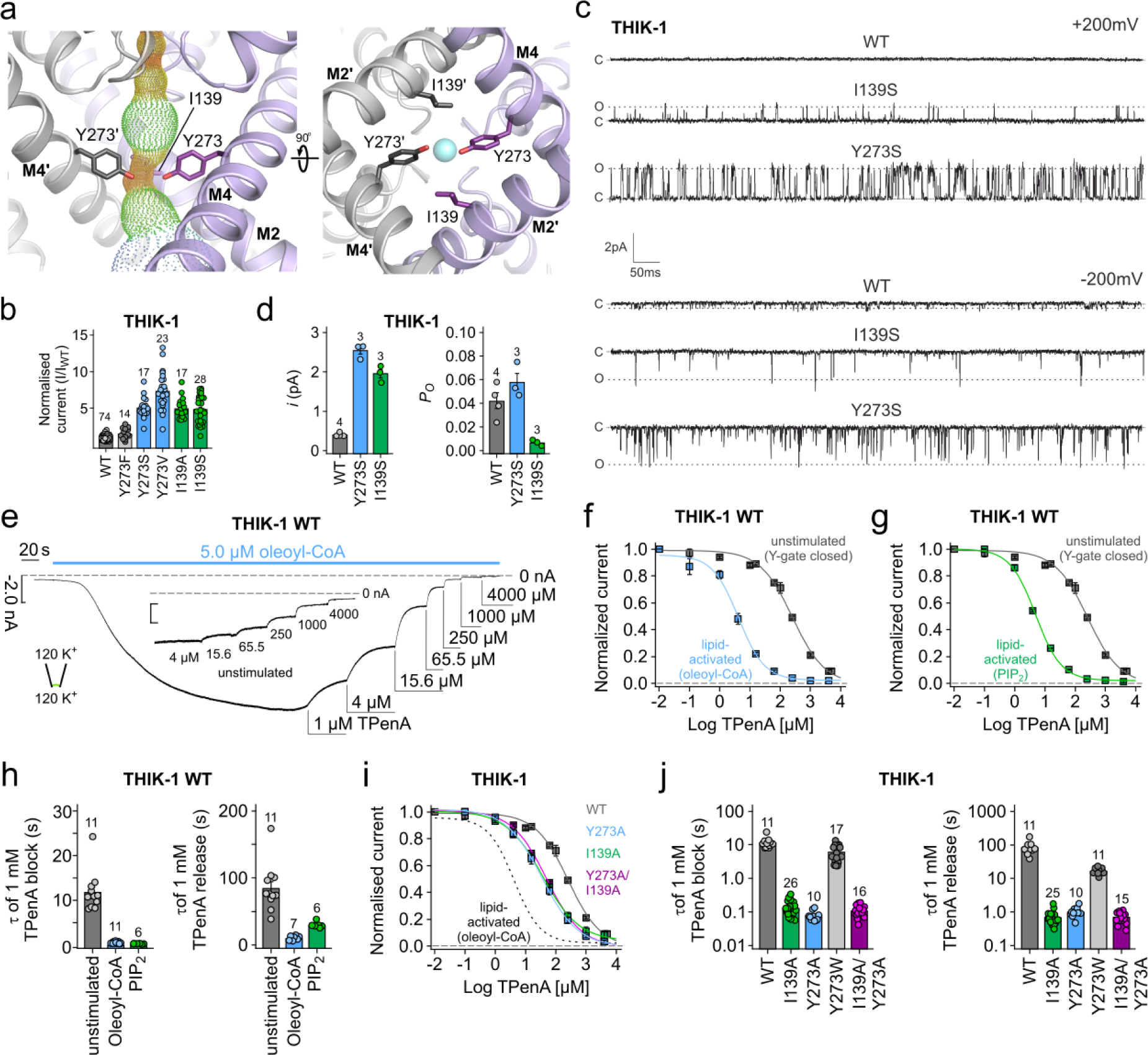
THIK-1 Y-gate regulated by lipids. **a,** The Y-gate viewed from the side (M2’ hidden for clarity), and from the bottom, showing residue I139 in the same horizontal level as Y273 and also part of the constriction formed by the Y-gate. **b,** Relative whole-cell current amplitudes of WT THIK-1 and channels with mutations in the Y-gate. All currents are normalised to WT. **c,** Cell-attached recordings of 1s duration at ± 200mV containing single WT, I139S and Y273S THIK-1 channels, as indicated. The closed (c) and open channel (o) levels are indicated. **d,** Comparison of single-channel open probability (*P_o_*) and single channel current amplitude (*i*) of WT, I139S and Y273S THIK-1 channels in cell-attached patches at −200mV. Numbers above the bars denote the number of experiments. **e,** Representative recording at −80 mV from an inside-out patch containing WT THIK-1 channels with symmetrical K^+^ concentrations (120 mM [K^+^]) at pH 7.4. Channel currents were activated with 5 µM oleoyl-CoA) applied to the intracellular side of the membrane and then inhibited dose-dependently with TPenA as indicated. Inlay shows equal inhibition with TPenA at the unstimulated basal state of the channels. **f,g,** Analysis of the apparent affinity for TPenA in the indicated states from recordings in panel e showing increased TPenA sensitivity after lipid activation. **h,** Analysis of TPenA kinetics for block and release of WT THIK-1 in unstimulated and lipid-activated states. **i**-**j,** Analysis of apparent TPenA affinity (**i**) and kinetics (**j**) for WT and indicated THIK-1 mutants from recordings as in the inlay in panel e. All values are given as mean ± s.e.m with number (n) of individual recordings indicated above the bars.

To examine the functional significance of this constriction we measured the activity of mutations in this motif (Y273S and I139S). Following expression in *Xenopus* oocytes, we found both mutants produced markedly increased whole-cell K^+^ currents compared to WT THIK-1, thus indicating an important role for these residues in the control of channel activity (**Figure 2b**).

We therefore examined their effects in more detail at the single channel level. Consistent with previous studies^18,19^, detection of single channel activity for WT THIK-1 in cell-attached patches with symmetrical K^+^ (140 mM) proved to be challenging. Small flickery openings were reliably resolved only at very negative potentials at (−200 mV) but not at positive potentials (up to +200mV) (**Figure 2c**). This suggest that at positive potentials the openings are either so fast they are filtered out, or that their individual conductance is so low that it is beyond the resolution of our recording system.

In contrast, both the Y273S and I139S mutants produced a marked increase in the single-channel conductance (∼6.5-fold greater than WT for Y273S, and ∼5-fold greater for I139S at −200 mV), as well as detectable openings at positive membrane potentials (**Figure 2c-d and Supplementary Figure 2a**). Interestingly, in comparison to WT THIK-1 at −200 mV, Y273S had no obvious effect on channel open-probability (*P_O_*), whilst the *P_O_* of I139S appears reduced due to an apparent increase in the stability of long closed states (**Figure 2c-d**).

We next examined the effect of a more conservative substitution at this position; the Y273F mutation only produced ∼1.5-fold increase in whole-cell currents (**Figure 2b**) with single channel currents indistinguishable from WT THIK-1 at −200mV (*i* = 0.47 ± 0.09 pA (n=3), *P_o_* = 0.058 ± 0.04, n=3). By contrast, the Y273V mutation produced >7-fold increase in whole-current (**Figure 2b**) that was also matched by a substantial increase in single channel conductance (*i* =-2.75 ± 0.01 pA at −200mV; n=3) (**Supplementary Figure 2a**).

Overall, this demonstrates that the effects of these mutations are complex and affect both K^+^ permeation and the dynamics of gating (i.e., *P_o_*). The Y273 side-chain therefore plays an important role in both these processes, and we hereafter refer to this structural motif as the Y-gate.

### The Y-gate opens upon lipid activation

If this structural motif is part of a physiologically-relevant gating mechanism, then it should open and close in response to regulatory inputs. It has previously been shown that THIK-1 is directly activated by polyanionic lipids such as PIP_2_ and oleoyl-CoA^17^. We therefore examined whether lipid regulation directly influences the Y-gate by measuring accessibility of pore blockers to the inner cavity above the Y-gate.

Like most K_2P_ channels, THIK-1 can be inhibited by intracellular application of tetrapentylammonium (TPenA). Consistent with this, we found that intracellular application of TPenA to excised patches from oocytes expressing THIK-1 produced inhibition with an IC_50_ of 225 ± 13 µM (n = 12) and that mutation of key residues immediately below the filter within the predicted consensus binding site for TPenA (T110A, C135A, T237A and V269A) reduced this inhibition (IC_50_ = 1284 ± 174 (n = 11), 1063 ± 140 (n = 8), 751 ± 69 (n = 10), 2203 ± 472 µM (n = 10), respectively) (**Supplementary Figure 2b-c**). This confirmed that, like other K2P channels, THIK-1 exhibits direct pore block by TPenA just below the SF. Strikingly, the kinetics of TPenA block and release were the slowest of all the K2P channels examined so far (τ_block_ = 12 ± 1 s and τ_release_ = 85 ± 11 s; n = 11) (**Supplementary Figure 2d**) consistent with the fact that the Y-gate clearly restricts access of TPenA to its binding site below the SF.

To address whether lipid activation regulates the Y-gate, we next measured the properties of TPenA inhibition in lipid-activated THIK-1 WT channels and observed that TPenA sensitivity was markedly increased by both oleoyl-CoA activation (IC_50_ = 3.9 ± 0.3; n = 22) and by 10 µM PIP_2_ activation (IC_50_ = 4.6 ± 0.2; n = 5) (**Figure 2e-g**). Furthermore, both the kinetics and efficacy of TPenA block were increased by lipid activation (oleoyl-CoA τ_block_ = 0.5 ± 0.1 s, τ_release_ = 11 ± 1 s; PIP_2_ τ_block_ = 0.31 ± 0.02 s and τ_release_ = 30 ± 2 s; n ≥ 6; WT inhibition with 15.6 µM = 11 ± 1 %, in oleoyl-CoA 78 ± 1 % and in PIP_2_ 74 ± 1 %, n ≥ 5, respectively) (**Figure 2h**). In addition, the sensitivity of TPenA inhibition was increased by activatory mutations within the Y-gate (Y273A: IC_50_ = 38 ± 7; n = 6; I139A IC_50_ = 38 ± 3; n = 13) as were the kinetics of TPenA inhibition. As a control, substitution with a bulky aromatic (Y273W) had little or no effect on these kinetics (**Figure 2i-j**). These results therefore demonstrate that anionic lipid activation dynamically changes the access of TPenA to its binding site by opening of the Y-gate.

The residues which comprise the Y-gate are also conserved in the related THIK-2 channel. Using an N-terminal ER-retention mutant (THIK-2*) which traffics to the membrane^20^, we found that mutation of the Y-gate (Y292A) in THIK-2* also produced a gain-of-function that affected both the sensitivity and kinetics of TPenA inhibition (**Supplementary Figure 2 e-g**).

Furthermore, it was recently shown that THIK-1 channels are involved in apoptotic processes such as cell shrinkage, via caspase-8 (CASP8) that cleaves the C-terminus of THIK-1 ^21^. Interestingly, truncation of the channel at this CASP8 cleavage site (G331x) also resulted in a gain-of-function and an increased TPenA sensitivity (G331x IC_50_ = 79 ± 17; n = 7) in comparison to WT (**Supplementary Figure 2h**), suggesting that CASP8 cleavage increases channel activity by promoting opening of the Y-gate. Together these results clearly demonstrate that this Y-gate motif integrates a variety of physiologically-relevant signals into a gating process that is conserved in this subfamily of K2P channels.

### Halothane inhibition also acts via the Y-gate

THIK-1 derives its original name from its sensitivity to the volatile anaesthetic, halothane. However, the binding site for halothane and its mechanism of inhibition remain unknown. To examine whether halothane inhibition also operates via the Y-gate we first tested for any direct competition with TPenA and found that halothane inhibition was reduced in the presence of TPenA (**Figure 3a-b**) indicating that it may also bind within the inner cavity.

**Figure. 3.**
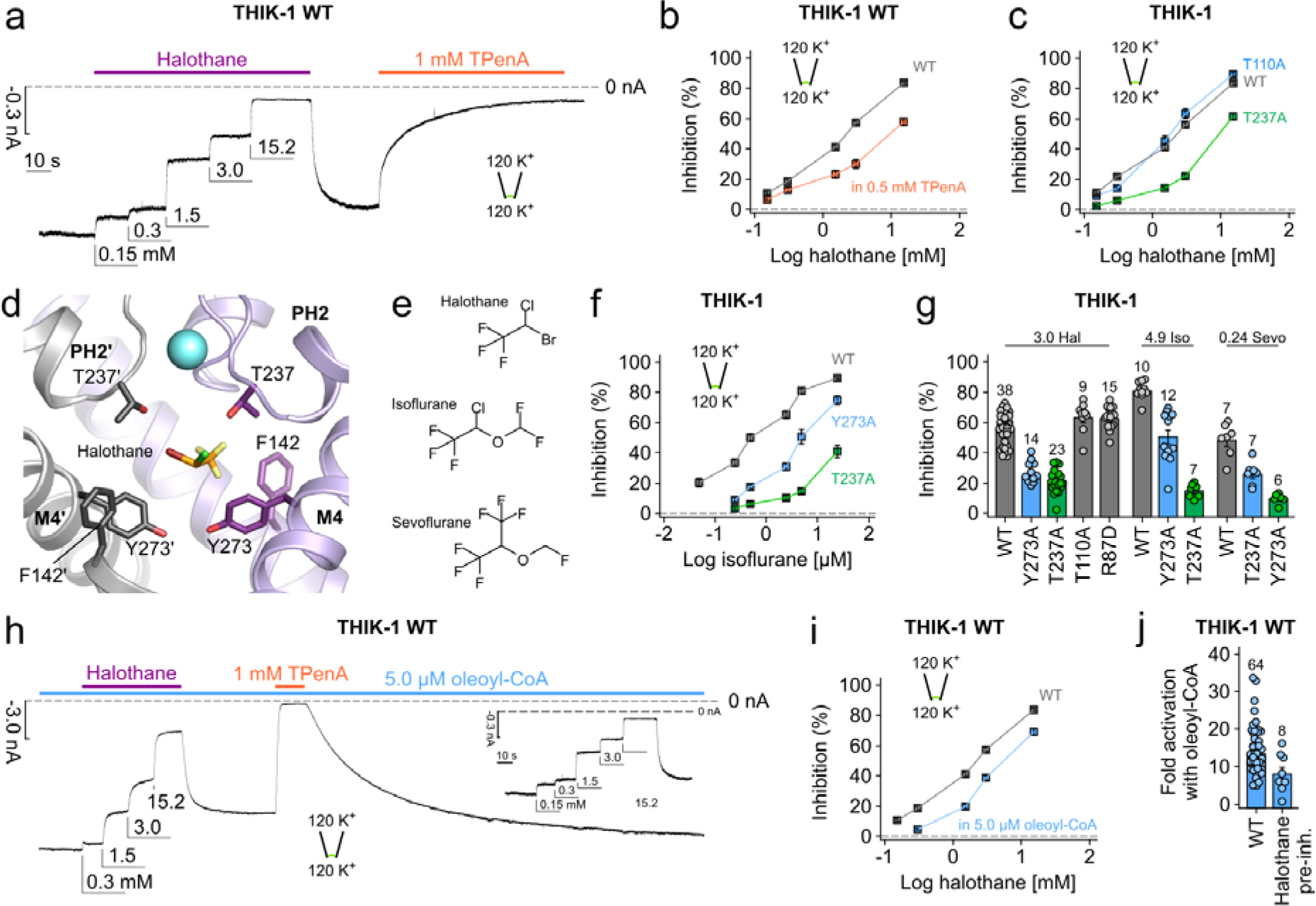
Inhibition by volatile anaesthetics involves both the filter and Y-gate. **a,** Representative recording at −80 mV from an inside-out patch containing WT THIK-1 channels with symmetrical K^+^ concentrations (120 mM [K^+^]_ex._/120 mM [K^+^]_int._) at pH 7.4. Channel currents were inhibited dose-dependently with increasing concentrations of halothane applied to the intracellular side of the membrane. Note, halothane effects can be washed and recovered and the currents inhibited with TPenA. **b,** Analysis of halothane inhibition for THIK-1 WT from recordings as in panel a in the absence (gray) and presence (orange) of 0.5 mM TPenA which produces ∼ 80 % block of initial currents. **c,** Analysis of halothane inhibition from recordings as in panel a for WT THIK-1 and indicated mutant channels. **d,** THIK-1 with docked halothane in the vestibule. Residues in close proximity are highlighted as sticks. For clarity, residues 121-138 in M2’ as well as PH2’ are not shown. **e,** Comparison of the structures of halothane, isoflurane and sevoflurane. **f,** Analysis of isoflurane inhibition of WT THIK-1 and channel mutants (i.e., Y273A and T237A). **g,** Summary of volatile anaesthetic inhibition with either 3.0 mM halothane, 4.9 mM isoflurane and 0.24 mM sevoflurane for WT THIK-1 and mutant channels as indicated. **h,** Representative recording under conditions as in panel a showing dose-dependent halothane inhibition for THIK-1 channels activated with 5.0 µM oleoyl-CoA. **i**, Analysis of halothane inhibition in the absence and presence of 5.0 µM oleoyl-CoA from recordings as in panels a and h. **j,** Fold activation of WT THIK-1 with 5.0 µM oleoyl-CoA in the absence and presence of 15.2 mM halothane. Values are given as mean ± s.e.m with number (n) of individual recordings indicated above the bars.

In other K2P channels, TPenA interacts with the conserved threonines of the **T**xGYG motif at the entrance to the filter (T110 in SF1 and T237 in SF2). Interestingly, mutation of either of these two threonines reduced TPenA inhibition (**Supplementary Figure S2b**), but only the T237A mutation in SF2 reduced halothane inhibition (**Figure 3c**), thus suggesting an off-centre, lateral binding site within this asymmetric cavity. It also suggests that halothane may not directly block the pore of THIK-1. Consistent with this, changes in extracellular K^+^ concentration had no effect on the efficacy of halothane inhibition of a more open, Y-gate gain-of-function mutant THIK-1 at −120 mV, indicating no knock-off effect occurs and that open channel block by halothane is unlikely (**Supplementary Figure 3a**).

Using the cryoEM structure of THIK-1 we next performed docking studies and identified a potential halothane binding site on one side within the inner cavity between F142 on M2, Y273 on M4, and T237 in SF2 (**Figure 3d** and **Supplementary Figure 3b**). In support of this, mutations within this predicted binding site affected inhibition by halothane as well the structurally related isoflurane and sevoflurane (**Figure 3e-g and Supplementary Figure 3c**). Docking also revealed similar binding sites for both isoflurane and sevoflurane, suggesting a conserved binding site for these volatile anaesthetics within the inner cavity (**Supplementary Figure 3e-f**).

Furthermore, we found that opening of the Y-gate by lipid activation, reduced both halothane and isoflurane inhibition (**Figure 3h, i and Supplementary Figure 3d**), and that pre-inhibition of WT THIK-1 channels with halothane reduced the extent of activation by 5 µM oleoyl-CoA (14 ± 1-fold, n = 64 for WT, *vs*. 8 ± 2-fold, n = 8, in the presence of halothane) (**Figure 3i**). Overall, these results indicate that halothane binding within the inner cavity also dynamically regulates opening and closing of the Y-gate.

### Extracellular ion exit pathway

The extracellular cap domain above the filter results in a bifurcated exit route for K^+^. Mutations within these pathways in other K2P channels have been shown to influence channel properties, in particular their sensitivity to blockers that interact with the ‘Keystone Inhibitor Site’ (KIS) immediately above the filter^22^, and obstruction of the pathway by nanobodies has also been shown to inhibit channel activity^23^.

In most K2Ps it is the presence or absence of negative charges at the KIS which affects channel conductance and pharmacology^24,25^. By marked contrast, the structure of THIK-1 reveals an unusually large number of positive charges, mostly arginine residues, distributed throughout the cap domain, several of which sit directly above the filter within the KIS (**Figure 1e** and **Supplementary Figure 4a**). In particular, the R83 and R87 side chains from each subunit position four positive charges directly above the filter and S0 K^+^ binding site in a manner likely to influence K^+^ permeation through the filter itself and also through the extracellular exit pathways that they line (**Figure 4a**).

**Figure 4.**
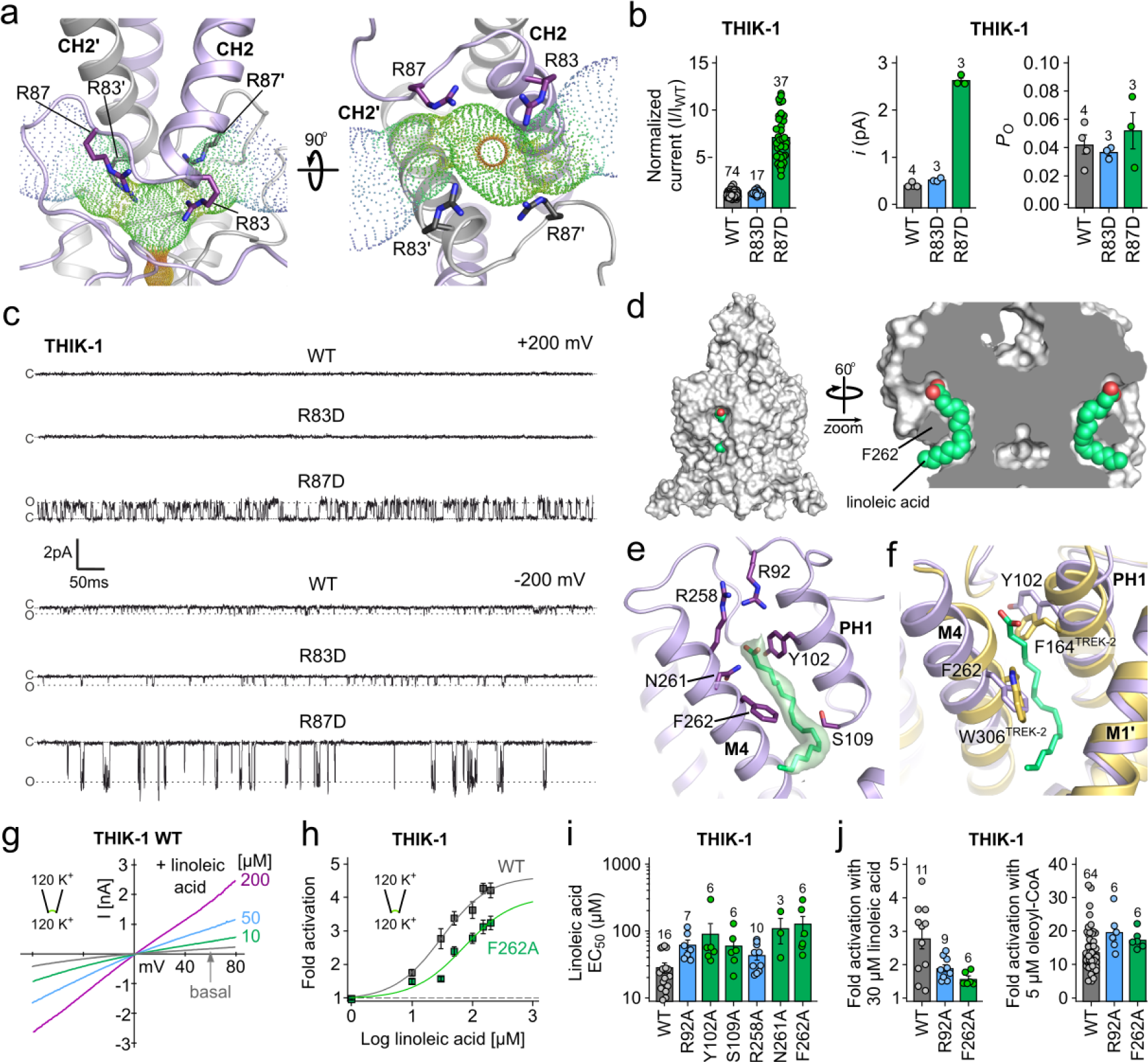
Regulation of THIK-1 activity by charged residues and lipids. **a,** Bifurcated extracellular ion exit pathway for K^+^ showing the orientation of positively-charged residues in that region (R83 and R87). **b,** Left: relative whole cell current amplitudes of WT and mutant THIK-1 channels (R83D and R87D). Right: Analysis of the single channel current amplitudes (*i*) and open probability (*P_O_*) from n ≥ 3 separate recordings as shown in panel c for WT and mutant THIK-1 channels in cell-attached patches at −200mV. **c,** Representative single channel recordings of WT THIK-1 compared to R83D and R87D mutant channels. Recordings shown at ± 200 mV in cell-attached configuration. The closed (c) and open (o) channel levels are shown. **d,** Surface representation of THIK-1 and cutaway showing two linoleic acid molecules (green) in the curved lipid binding pocket. **e,** The lipid binding pocket, with residues that have been mutated for functional studies highlighted as sticks. **f,** Structural overlay of THIK-1 (purple) with TREK-2 (yellow) showing position of F262 and Y102 in THIK-1 with equivalent W306 and F164 in TREK-2 which rotate during channel activation. **g,** Representative recording from an inside-out patch containing WT THIK-1 with symmetrical K^+^ concentrations (120 mM [K^+^]) at pH 7.4. Channel currents were activated with indicated concentrations of linoleic acid. **h,** Analysis of fold activations from recordings as in panel g for WT and F262A mutant channels. **i,** EC_50_ values from dose-response curves as in panel h for WT and mutant THIK-1. **j,** Fold activation of WT and R92A and F262A mutant THIK-1 channels, respectively with either 30 µM LA or 5 µM oleoyl-CoA. Values are given as mean ± s.e.m with number (n) of individual recordings indicated above the bars.

Interestingly, a charge reversal at the first of these positions (R83D) produced channels with whole cell currents ∼1.5-fold larger than WT THIK-1 (**Figure 4b**), but still with no resolvable single-channel openings at positive potentials, and only a modest effect at inward potentials (**Figure 4c**). However, a charge reversal at the second position (R87D) produced whole cell currents ∼7-fold larger than WT THIK-1, and single channel recordings revealed a markedly increased single-channel conductance at both positive and negative voltages (**Figure 4b-c).** The R87D mutant also exhibited kinetics that were distinct from channels with mutations in the Y-gate (Y273S), with openings clustered into longer bursts and clearly separated intra- and inter-burst closed states (**Supplementary Figure 4b**).

Also, in comparison to Y273S, the *P_O_* of R87D channels was steeply voltage-dependent at positive membrane potentials **(Supplementary Figure 4c,e)** suggesting a negative charge at this position not only stimulates K^+^ efflux through the filter, but also promotes channel opening (i.e., increases *P_O_*) via stabilising the open conformation of the SF. Furthermore, although the *P_O_* of the R87D and Y273S mutations approach similar values at +100mV, unlike Y273S, R87D single channels exhibit inward rectification **(Supplementary Figure 4d-e)**. This suggests that a negative charge at position 87 can also hinder the exit of K^+^ via extracellular ion exit pathways to reduce single-channel conductance at positive membrane potentials. Intriguingly, human microglia have a relatively positive resting membrane potential (−30 to −40 mV) that can be depolarised even further with selective THIK-1 inhibitors^5^. It will therefore be interesting to determine how these positive charges in THIK-1 contribute to the unusual electrical properties of these cells.

The well-behaved single channel behaviour of the R87D mutant also presents an opportunity to examine the effects of halothane inhibition in more detail as its effects on WT THIK-1 single channels would be difficult to measure. Excised patch experiments show comparable inhibition for WT and R87D mutant channels **(Supplementary Figure 5a,b)** and we found that halothane induces destabilisation (shortening) of both open and closed states with the former driving the inhibitory effect. **(Supplementary Figure 5c,d)**.

### A lipid bound within the K2P modulator pocket

A cryptic small molecule binding site has previously been identified in structures of TREK channels in complex with certain activators (ML335 and ML402)^26^. This pocket is located at the PH1-M4 interface and modulation of this interface dynamically regulates K2P channel activity through direct effects on the filter-gate^27^. It was therefore interesting to observe density within this pocket consistent with a fatty acid (**Figure 1f**, **Figure 4d-f and Supplementary Figure 1e**).

Because this binding pocket is also highly curved the bound lipid is likely to be a polyunsaturated fatty acid (PUFA) and we found that the Omega-3 essential fatty acid, linoleic acid (18:2) fits well into this density (**Figure 4d-f**). Two arginine side chains (R92 and R258) are in close proximity to the headgroup of this lipid and F262 on M4 contacts the middle of the lipid (**Figure 4e**). This is of particular interest because in many other K2P channels, F262 is a conserved tryptophan predicted to rotate inwards into this pocket during channel activation^23,28^ (**Figure 4f**). Any lipid bound within this site might therefore modulate THIK-1 activity and thus influence microglial function.

Interestingly, we found that linoleic acid directly activates THIK-1 in excised patches with an EC_50_ 29 ± 5 µM, n = 16 (**Figure 4g**), and that mutations within this pocket reduce both the extent and affinity of this activation (**Figure 4h-i**). In support of this as a unique binding site for this PUFA, these mutations do not affect oleoyl-CoA activation. It is also unclear how such anionic lipids with larger headgroups could be accommodated within this pocket. Consistent with this, truncation of the C-terminus (G331x at the CASP8 cleavage site) reduced anionic lipid activation (**Supplementary Figure S5e**) highlighting these two separate and distinct sites for lipid activation of THIK-1. Thus, although the precise identity of the lipid bound within this modulator pocket remains to be determined unambiguously, this curved site can only accommodate a restricted range of PUFAs, with linoleic acid the prime candidate. This is supported by the activatory effect of this lipid and of mutations within this binding pocket. The potential role of this lipid in regulating THIK-1 therefore deserves further investigation because microglia can sense a variety of extracellular lipids that stimulate their inflammatory response and changes in linoleic acid have been shown to influence microglial activity^29,30^.

Overall, this study provides structural insight into the gating and permeation of this unusual subfamily of K2P channels, and reveals a unique lower cytoplasmic Y-gate that responds to physiologically-relevant signals. Furthermore, by providing a structural framework for the optimisation of both THIK-1 inhibitors and possible activators, our results have major implications for the modulation of microglial function and future therapeutic strategies for the treatment of neurodegenerative disorders.

## Methods

### Cloning and protein expression

Human *KCNK13*, encoding residues Gly9 to Gly297 was subcloned into a modified pFastBac vector, encoding a HRV 3C protease site, a decahistidine tag and a FLAG tag, using ligase-independent cloning. Bacmid DNA was generated using the Bac-to-Bac system and the resulting bacmid transfected into *Spodoptera frugiperda* (Sf) 9 insect cells. The resulting virus was amplified twice and used for large scale infections of Sf9 cells at a density of 2 × 10^6^ cells/mL, using 5% v/v virus per litre of cells, and grown for 72 h at 27°C. Cells were harvested by centrifugation at 900 x g, 10 min and stored at −80°C prior to protein purification.

### Purification

Cells were resuspended in 40 ml breaking buffer (50 mM HEPES pH 7.5, 200 mM KCl, 5% v/v glycerol) per litre of initial cell culture volume and lysed using an EmulsiFlex-C5 high-pressure homogenizer (Avestin), prior to solubilisation in 1% w/v octyl glucose neopentyl glycol (OGNG) and 0.1% w/v cholesteryl hemisuccinate (CHS) for 1 h. Residual debris was pelleted by centrifugation at 35,000 × g, for 1 h. To collect the protein, the supernatant was incubated with 1 ml 50% v/v Talon resin (Clontech) and 5 mM imidazole pH 8.0 per initial litre of cell culture for 1 h. The resin was collected and washed with 30 column volumes of wash buffer (50 mM HEPES pH 7.5, 200 mM KCl, 5% v/v glycerol, 20 mM imidazole pH 8.0, 0.18% w/v OGNG, 0.018% w/v CHS) and the final volume of resin and buffer was adjusted to 2 column volumes. On-column cleavage and deglycosylation of the protein was done by adding 150 µg hexahistidine tagged HRV 3C protease and 50 µg hexahistidine tagged PNGaseF, prior to incubation overnight. The flow-through was collected and concentrated to 500 µl and subjected to size exclusion chromatography on a Superose 6 Increase 100/300 GL column (Cytiva) in gel filtration buffer (20 mM HEPES pH 7.5, 200 mM KCl, 0.12% w/v OGNG, 0.012% w/v CHS). All purification steps were performed on ice or at 4°C.

### CryoEM grid preparation and data collection

Electron microscopy was provided through the Central Oxford Structural Molecular Imaging Centre (COSMIC) facility. Grids were prepared by plasma treating holey gold grids (GF-1.2/1.3, 300 mesh, 45 nm film) and adsorbing 3 µl THIK-1 at 4 mg/ml, blotted for 3-6 s at 100% humidity at 4°C, and vitrified in liquid ethane, using a Vitrobot Mark IV (Thermo Fischer Sceintific). Data were collected on a Titan Krios G3 (FEI) with a K3 direct detection camera (Gatan) and a BioQuanum imaging filter (Gatan), at 300 kV in counted superresolution bin 2 mode and 105,000 × magnification, with a pixel size of 0.832 Å, and a total dose of 42.54 e^−^/Å^2^ over 40 fractions. A total of 21,171 movies were collected over a defocus range of −1.0 to −2.3 µm.

### Data processing

Motion correction and contrast transfer function (CTF) estimation was done on a subset of 1060 movies, using the preprocess_stream pipeline within SIMPLE^31^. Manual particle picking, followed by autopicking, extracting and one round of 2D classification in SIMPLE. Particles belonging to the accepted 2D classes were imported into cryoSPARC^32^. All further processing steps were done in cryoSPARC, unless stated otherwise. The particles were subjected to two rounds of 2D classification and one round of *ab initio* reconstruction with five classes, and templates were generated from one of the classes. The templates were then used for repicking of the micrograph subset. The extracted particles were subjected to similar steps to generate improved templates.

The full dataset was motion corrected in SIMPLE, using the preprocess_stream pipeline, and motion corrected micrographs were imported into cryoSPARC where they underwent patch CTF estimation. Of the initial 21,171 micrographs, 18,838 were kept and particles then picked from these using the previously generated templates. A total of 9,257,778 particles were extracted with a box size of 256 pixels, and underwent one round of unmasked 2D classification and five rounds of 2D classifications with a 125 Å spherical mask. The selected 1,240,628 particles were subjected to an *ab initio* volume reconstruction with five classes, applying no symmetry, followed by heterogenous refinement. Three of the classes, 878,207 particles in total, were further cleaned by one round of 2D classification, narrowing the set down to 765,828 particles. These were used for another round of ab initio reconstruction with five classes, applying C2 symmetry, followed by heterogenous refinement. The particles in one class (327,614), were further refined with homogenous and non-uniform refinement, applying C2 symmetry. They were then exported to RELION-3, using the csparc2star.py script within UCSF pyem (https://zenodo.org/records/3576630), and underwent Bayesian polishing^33^. After a final round of 2D classification, the cleaned and polished particle set (302,189) were used for non-uniform refinement with C2 symmetry applied. A final resolution of 3.16 Å was estimated with gold-standard Fourier shell correlations (FSCs) using the 0.143 criteria. The map was subsequently sharpened with a B-factor of −140.

### Model building and refinement

An initial THIK-1 model was built manually in Coot^34^. An elongated density was observed behind the filter, and we chose to model linoleic acid, in this density, based on the fit to the map. The model was refined with *phenix.real_space_refine*^35,36^, with NCS constraints and restraints for the linoleic acid generated using the Grade2 server, and otherwise default settings. The refined model was further improved in ISOLDE^37^, within ChimeraX^38^, and used to generate a C2 symmetric model which was subjected to real space refinement, with NCS constraints and ligand restraints as before, but without rotamer or Ramachandran restraints and restraining the input model to the ISOLDE generated A chain. This model was then subjected to a final round of adp refinement. The radii of the channel pore and the extracellular ion pathways were calculated using HOLE^39^.

### Molecular docking

Molecular docking using was used with this structure of THIK-1 to determine and evaluate the most probable binding poses for halothane, isoflurane and sevoflurane. Based on the result of the competition assay of halothane with TPenA, it is expected that halothane binds within the pore cavity close to the known binding sites for QA blockers. Molecular docking was therefore first used to explore possible binding poses of halothane within the structure with the potassium density at the filter S4 site defined as the center of the docking pocket. A receptor grid was calculated for THIK-1 with a box size of 10 x 10 x 10 Å. Docking of halothane, isoflurane and sevoflurane into the pocket was subsequently carried out. The 3 docking poses with the highest scores were selected and further validated by electrophysiological analysis.

### Molecular biology

Human K_2P_2.1 THIK-1 (GenBank accession number: NM_022054) and human K_2P_10.1 THIK-2 (NM_022055.1) were used in this study. For K^+^ channel constructs expressed in *Xenopus* oocytes the respective K^+^ channel subtype coding sequences were subcloned into the oocyte expression vector pSGEM or pFAW^40^ and verified by sequencing. All mutant channels were obtained by site-directed mutagenesis and verified by sequencing. To increase surface expression and macroscopic currents, measurements of THIK-2 channels used a mutated ER retention motif, (R11A/R12A/R14A/R15A/R16A; THIK-2*)^20^. Vector DNA was linearized with NheI or MluI and mRNA synthesized *in vitro* using the SP6 or T7 AmpliCap Max High Yield Message Maker Kit (Cellscript, USA) or HiScribe® T7 ARCA mRNA Kit (New England Biolabs).

### Two-electrode voltage-clamp (TEVC) measurements

Electrophysiological studies were performed using the TEVC technique in *Xenopus* oocytes. All animal use conformed to the guide for the Care and Use of laboratory Animals (NIH Publication 85-23) and all. experiments using *Xenopus* toads were approved by the local ethics panels. Oocytes were stored at 17 °C in ND96 recording solution (in mM): 96 NaCl, 2 KCl, 1.8 CaCl_2_, 1 MgCl_2_, 5 HEPES (pH 7.5 adjusted with NaOH/HCl) supplemented with Na-pyruvate (275 mg/l), theophylline (90 mg/l), and gentamicin (50 mg/l). Oocytes were injected with 1 ng of mRNA for WT or mutant channels and incubated for 24-48hrs days at 17°C. Two-Electrode Voltage Clamp recordings then performed as previously described ^41,42^.

### Macroscopic patch-clamp measurements

Oocytes were obtained as described above incubated at 17°C in a solution containing (mM): 54 NaCl, 30 KCl, 2.4 NaHCO_3_, 0.82 MgSO_4_ x7 H_2_O, 0.41 CaCl_2_, 0.33 Ca(NO_3_)_2_ x4 H_2_O and 7.5 TRIS (pH 7.4 adjusted with NaOH/HCl) for 1-4 days before use. Excised patch recordings in inside-out configuration conditions were performed at room temperature. Patch pipettes were made from thick-walled borosilicate glass GB 200TF-8P (Science Products, Germany), had resistances of 0.2 - 0.5 MΩ (tip diameter of 10 – 25 µm) and filled with a pipette solution (in mM): 120 KCl, 10 HEPES and 3.6 CaCl_2_ (pH 7.4 adjusted with KOH/HCl). Intracellular bath solutions and compounds were applied to the cytoplasmic side of excised patches for the various K^+^ channels via a gravity flow multi-barrel pipette system. Intracellular solution had the following composition (in mM): 120 KCl, 10 HEPES, 2 EGTA and 1 Pyrophosphate (pH adjusted with KOH/HCl). Currents were recorded with an EPC10 amplifier (HEKA electronics, Germany) and sampled at 10 kHz or higher and filtered with 3 kHz (−3 dB) or higher as appropriate for sampling rate.

### Single channel patch-clamp measurements

Single channel currents were recorded with Axopatch 200B amplifier via a Digidata 1440A digitizer (Molecular Devices). Data were filtered at 2kHz and recorded at a 200-kHz sampling rate with program Clampex (Molecular Devices). Pipette solution and bath solution for cell-attached experiments contained (in mM): 140 KCl, 2 MgCl_2_, 1 CaCl_2_, 10 HEPES (pH 7.4 adjusted with KOH/HCl). For inside-out experiments, the bath solution contained (in mM): 140 KCl, 2 MgCl_2_, 1 CaCl_2_, 10 HEPES (pH 7.2 adjusted with KOH/HCl). All experiments were conducted at room temperature.

### Clinical drugs, chemical compounds and lipids

Tetrapentylammonium (TPenA), linoleic acid, L-α-PI(4,5)P_2_ (brain PI(4,5)P_2_, PIP_2_) (Sigma-Aldrich/Merck, Germany) and oleoyl-CoA (LC-CoA 18:1) (Avanti Polar Lipids, USA) were prepared as stocks (1 - 100 mM) in DMSO, stored at −80 °C and diluted to the final concentration in the intracellular recording solution.Halothane 99 % (Sigma-Aldrich/Merck, Germany), isoflurane (Baxter, Germany) and sevoflurane 100 % (AbbVie, Germany), respectively were added to the intracellular recording solution, shaken for 3 min. and after clear phase separation (∼ 5 min.) the intracellular solution was used for experiments within 15 min.

### Data acquisition and statistical analysis

Data analysis and statistics for macroscopic measurements were done using Fitmaster (HEKA electronics, version: v2×90, Germany), Microsoft Excel 2021 (Microsoft Corporation, USA) and Igor Pro 9 software (WaveMetrics Inc., USA). Recorded currents were analyzed from membrane patches at a voltage defined in the respective figure legend or with a voltage protocol as indicated in the respective figure. The fold activation (fold change in current amplitude) of a ligand (clinical drug, compound or lipid) was calculated from the following equation:

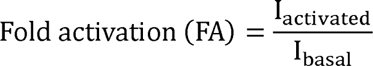

where I_activated_ represents the stable current level in the presence of a given concentration of a respective ligand and I_basal_ the measured current before ligand application. Percentage inhibition upon blocker application for a ligand was calculated from stable currents of excised membrane patches using the following equation below:

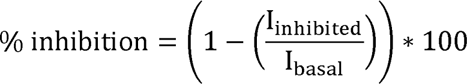

where I_inhibited_ refers to the stable current level recorded in the presence of a given concentration of the ligand and I_basal_ to the measured current before ligand application. The macroscopic half-maximal concentration-inhibition relationship of a ligand was obtained using a Hill-fit for dose-response curves as depicted below:

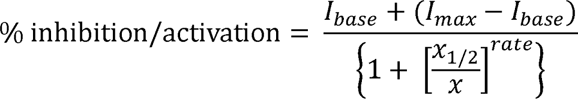

where base and max are the currents in the absence and presence of a respective ligand, x is the concentration of the ligand, x_1/2_ is the ligand concentration at which the activatory or inhibitory effect is half-maximal, rate is the Hill coefficient.

For analysis of block and release from block time constants (τ) current traces were fitted with a mono-exponential equation as depicted below:

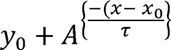

Data from individual measurements were normalized and fitted independently to facilitate averaging. Error bars in all figures represent the s.e.m. with numbers (n) above indicating the definite number of individual executed experiments. A Kolmogorow-Smirnow test was used to determine whether measurements were normally distributed. Statistical significance between two groups (respective datasets) was validated using an unpaired two-tailed Student’s *t*-test. Asterisks indicate the following significance: * *P* ≤ 0.05, ** *P* ≤ 0.01 and *** *P* ≤ 0.001.. Image processing and figure design was done using Igor Pro 9 (64 bit) (WaveMetrics, Inc., USA), PyMOL 2.4.1 (Schrödinger, LLC) and Canvas X Draw (Version 20 Build 544) (ACD Systems, Canada). For analysis of single-channels, recordings were idealized using 50% threshold criterion with Clampfit (Molecular Devices) at an imposed resolution of 50 µs. Analysis of amplitude and dwell-time distributions was performed in Origin (OriginLab Corporation). Critical time for burst analysis was determined using Magleby and Pallotta criterion^43^.

## Data Availability

All data within this study are included in the article and/or the Supplementary Information, and materials are available upon request. The cryoEM model and map of THIK-1 has been deposited in the Protein Data Bank and the EMDB database, respectively, under accession codes 9FT7, and EMD-50741.

## Supporting information

Higher Resolution Figures

## Acknowledgments

This work was directly supported by grants from the Biotechnology and Biological Sciences Research Council and Medical Research Council to S.J.T (BB/T002018/1, BB/S008608/1 and MR/W017741/1). It was also supported by the Wellcome Trust as part of the OXION Initiative (WT084655MA and 102161/B/13/Z). Further grants from the Deutsche Forschungsgemeinschaft supported the work of M.S. (SCHE 2112/1-2) and T.B (BA 1793/6-2) as part of the Research Unit FOR2518, DynIon. We thank Liz Carpenter and members of the Structural Genomics Consortium, as well as Alexander Baker and Anna Shaw for their contributions to the early stages of this project. We also thank the staff of the Central Oxford Structural Molecular Imaging Centre (COSMIC) facility for advice and assistance with sample preparation, screening and data collection setup, as well as with data processing and model refinement.

## Author contributions

S.J.T., K.E.J.R., M.S and P.P. conceived/designed the principal elements of the study. All authors (K.E.J.R, M.S., P.P, B.E., M.S.H., A.B, C.M., A.R., S.N., T.B, and S.J.T.) generated, analyzed or interpreted data, or generated materials. S.J.T, K.E.J.R and M.S. drafted the manuscript and all authors contributed to the final version.

## Competing interests

A.R is employed by Cerevance Ltd, Cambridge, UK.

## Notes

### Competing Interest Statement

AR is employed by Cerevance Ltd, Cambridge, UK.

