## Supplementary Information for "CryoEM Structure of the human THIK-1 K2P K^+^ Channel Reveals a Lower ‘Y-gate’ Regulated by Lipids and Anaesthetics"

### Supplementary Table 1

#### Cryo-EM data collection, refinement and validation statistics

| | THIK-1 (EMDB-50741) (PDB 9FT7) |
|---|---|
| **Data collection and processing** | |
| Magnification | 105,000 x |
| Voltage (kV) | 300 |
| Electron exposure (e–/Å$^2$) | 42.54 |
| Defocus range (μm) | -1 to -2.3 |
| Pixel size (Å) | 0.832 |
| Symmetry imposed | C2 |
| Initial particle images (no.) | 9,257,778 |
| Final particle images (no.) | 302,189 |
| Map resolution (Å) | 3.16 |
| FSC threshold | 0.143 |
| **Refinement** | |
| Model resolution (Å) | 3.16 |
| FSC threshold | 0.143 |
| Map sharpening *B* factor (Å$^2$) | -140 |
| Model composition | |
| Non-hydrogen atoms | 4016 |
| Protein residues | 508 |
| Potassium ion | 2 |
| Linoleic acid | 2 |
| *B* factors (Å$^2$) | |
| Protein | 64.73 |
| Potassium ion | 23.63 |
| Linoleic acid | 48.43 |
| R.m.s. deviations | |
| Bond lengths (Å) | 0.007 |
| Bond angles (°) | 1.064 |
| Validation | |
| MolProbity score | 0.85 |
| Clashscore | 1.26 |
| Poor rotamers (%) | 0 |
| Ramachandran plot | |
| Favored (%) | 98.82 |
| Allowed (%) | 1.18 |
| Disallowed (%) | 0 |

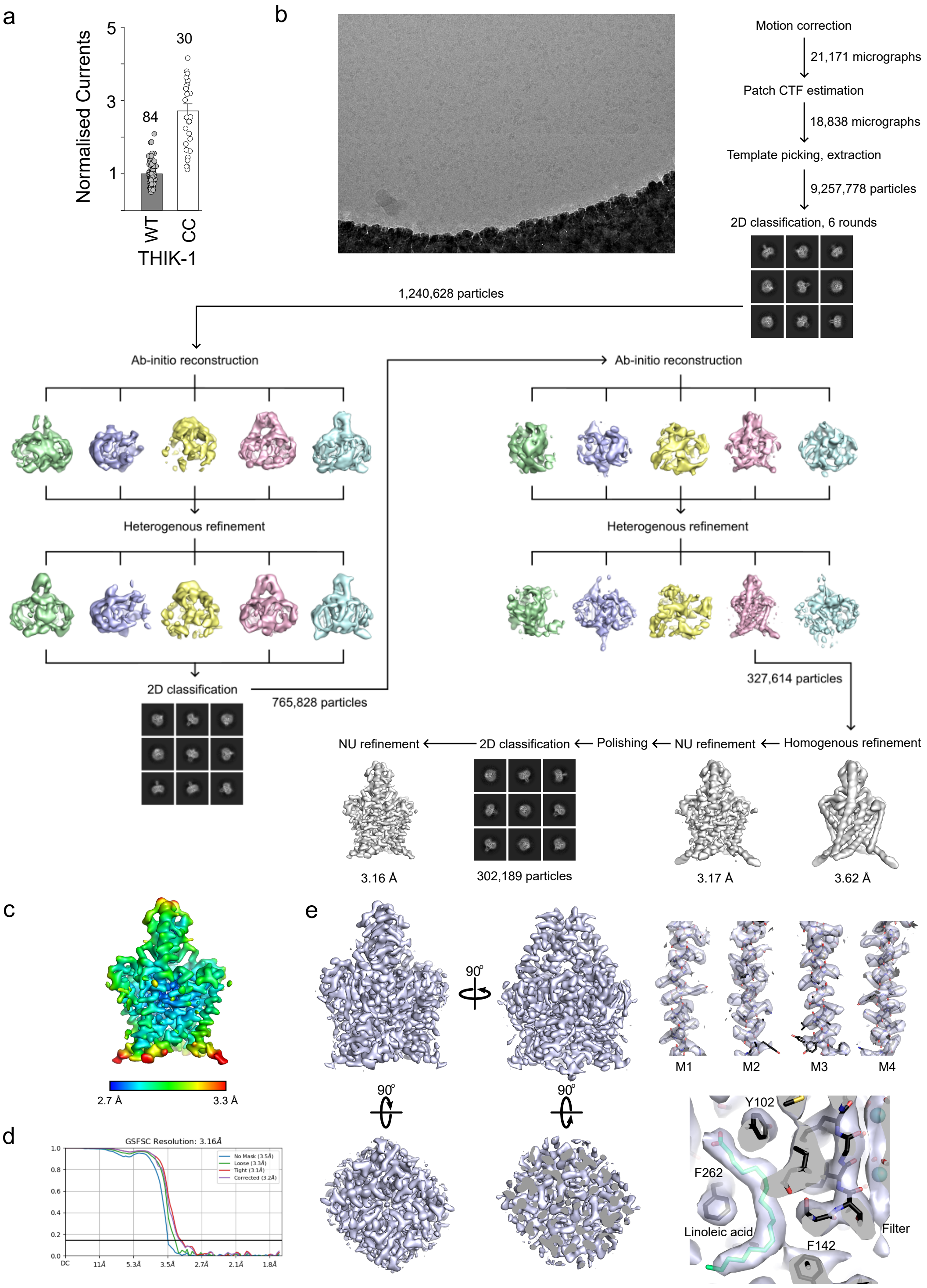

**Figure S1. Cryo-EM processing pipeline for THIK-1**. **a,** The cryoEM construct (CC) used for structural studies remains functionally active; whole-cell currents (recorded at +50 mV) are normalised to WT THIK-1. **b**, CryoEM imaging processing workflow. **c,** Local resolution estimation of the unsharpened map, determined within cryoSPARC. **d,** Gold standard FSC curve for resolution estimation, calculated within cryoSPARC. **e,** The sharpened map, overall views from the sides, bottom and slice through the top. Maps 2.6 Å around the M1-M4 helices and a close up view of the map in the lipid binding site.

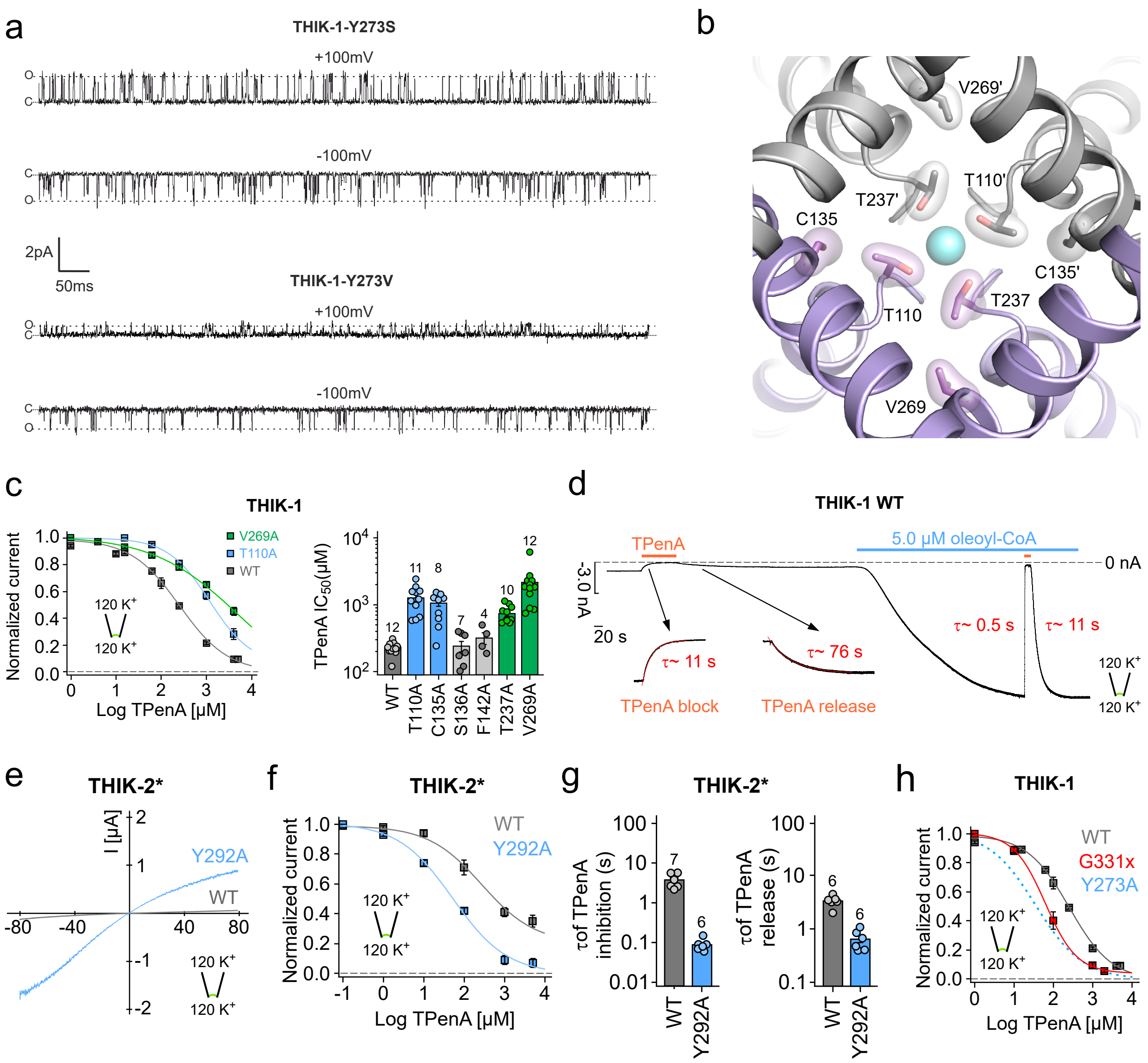

**Figure S2. Y-gating and TPenA inhibition in THIK channels**. **a,** Cell-attached recordings of 1 s duration at ± 100mV containing single Y273S and Y273V THIK 1 channels, as indicated. The closed (c) and open channel (o) levels are indicated. **b,** Bottom view towards the selectivity filter with residues involved in TPenA binding highlighted. The $K^+$ ion in the filter is represented in cyan. **c,** TPenA dose-response curves for WT (gray), T110A (blue) and V269 (green) mutant THIK-1 channels. TPenA $IC_{50}$ values for WT and indicated mutants. **d,** Representative recording at -80 mV from an inside out patch of *Xenopus* oocyte expressing WT THIK-1 channels with symmetrical $K^+$ concentrations (120 mM $[K^+]$ex./ 120 mM $[K^+]$int.) at pH 7.4. Channel currents were inhibited with 1 mM TPenA in the absence and presence of 5 µM oleoyl-CoA. **e,** Representative recording from inside out patches containing either THIK-2* (gray trace) or THIK-2* Y292A channels (blue trace) with symmetrical $[K^+]$ (120 mM $[K^+]$ex./120 mM $[K^+]$int.) at pH 7.4. **f,g,** TPenA dose-response curves (f) and block/release kinetics (g) for THIK-2* (gray) and THIK-2* Y292A (blue) mutant channels, respectively. **h,** TPenA dose-response curves for WT (gray) and G331x mutant THIK-1 channels. Note, dashed line indicates TPenA dose-response curve for THIK-1 Y273A channels. Values are given as mean ± s.e.m with number (n) of individual recordings indicated above the bars.

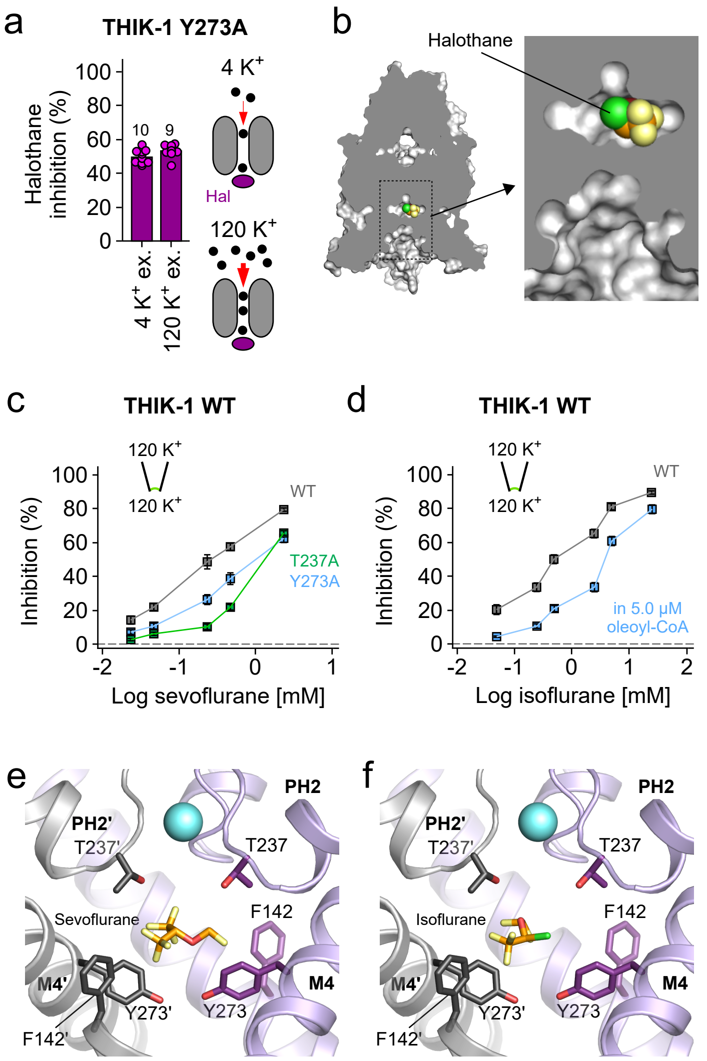

**Figure S3. Volatile anaesthetic inhibition in THIK-1 channels. a,** Analysis of halothane inhibition of THIK-1 channel currents recorded at -80 mV from excised patches at pH 7.4 with low (4 mM) and high (120 mM) extracellular $K^+$ indicating lack of effect of external [$K^+$] which would otherwise displace a pore blocker. **b,** Surface representation and expanded cutaway of THIK-1 showing halothane docked off centre within the inner cavity. **c,** Analysis of sevoflurane inhibition for WT (gray), THIK-1 T237A (green) and Y273A mutant channels (blue), respectively. **d,** Analysis of halothane inhibition in the absence and presence of 5.0 µM oleoyl-CoA. **e,f,** THIK-1 with docked sevoflurane (e) or isoflurane (f) in the inner vestibule. For clarity, residues 121-138 in M2’ and PH2’ are not shown. Values are given as mean ± s.e.m with number (n) of individual recordings indicated above the bars.

### Supplementary Figure S4

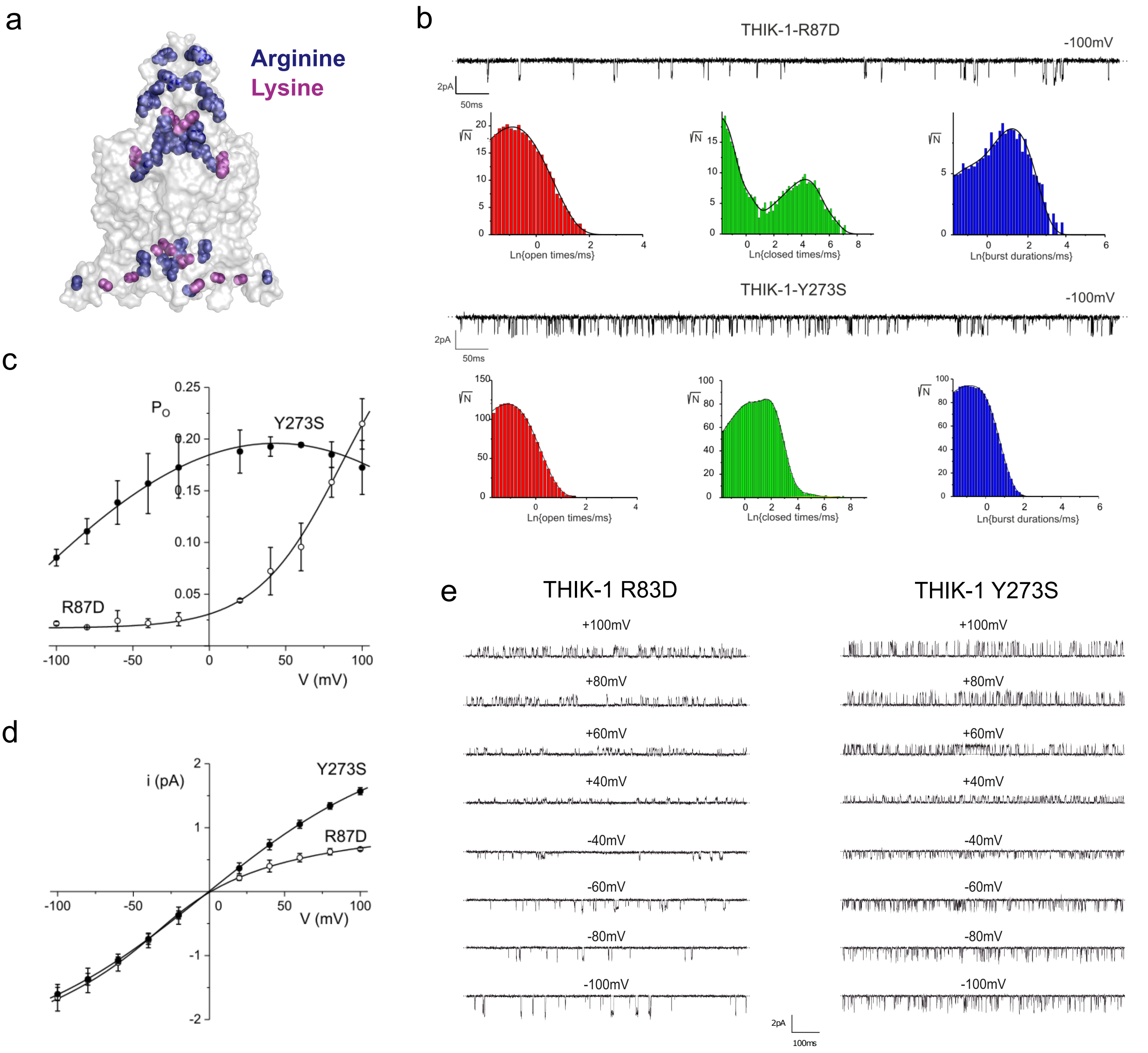

**Figure S4. Role of positive charges in the cap domain**. **a,** Transparent surface representation of THIK-1 showing the cluster of arginine (blue) and lysine (purple) residues in the cap domain and at the intracellular pore site. **b,** Comparison of single-channel kinetics of THIK-1-R87D (top) and THIK-1 Y273S (bottom) recorded in cell-attached configuration at -100 mV. Traces represent 1 s recordings with horizontal dotted lines indicating the zero current level. Below the traces are dwell time histograms of openings (red bars), closings (green bars) and bursts of openings (blue bars). The lines are probability density function fits to the data: $t_1$ = 0.31 ms, $A_1$ = 611; $t_2$ = 0.67 ms, $A_2$ = 529; (openings); $t_1$ = 0.14 ms $A_1$ = 902; $t_2$ = 0.58 ms, $A_2$ = 136; $t_3$ = 0.79 ms, $A_3$ = 11; $t_4$ = 14ms, $A_4$ = 13; $t_5$ = 66 ms, $A_5$ = 184; $t_6$ = 185 ms; $A_6$ = 42 (closings) and $t_1$ = 0.23 ms, $A_1$ = 36; $t_2$ = 3.5 ms, $A_2$ = 204 (bursts) for THIK-1 R87D, and $t_1$ = 0.17 ms $A_1$ = 1914; $t_2$ = 0.78 ms, $A_2$ = 9500; $t_3$ = 5.3 ms, $A_3$ = 17613; $t_4$ = 11 ms, $A_4$ = 1695; $t_5$ = 58 ms, $A_5$ = 37; $t_6$ = 300 ms; $A_6$ = 3 (closings) and $t_1$ = 0.12 ms, $A_1$ = 7900; $t_2$ = 0.59 ms, $A_2$ = 22976 (bursts) for THIK-1-Y273S. **c,** Comparison of voltage-dependence of single-channel open probability ($P_o$) and **d**, single-channel current amplitude (*i*) of THIK-1 R87D and THIK-1 Y273S mutant channels (n = 3 for all data points). Note, lines through the data are drawn by hand. **e,** Cell-attached recordings of 1 s duration at voltages between -100 mV and +100 mV of single THIK-1 R87D and THIK-1 Y273S, as indicated. The dotted lines represent zero current levels.

### Supplementary Figure S5

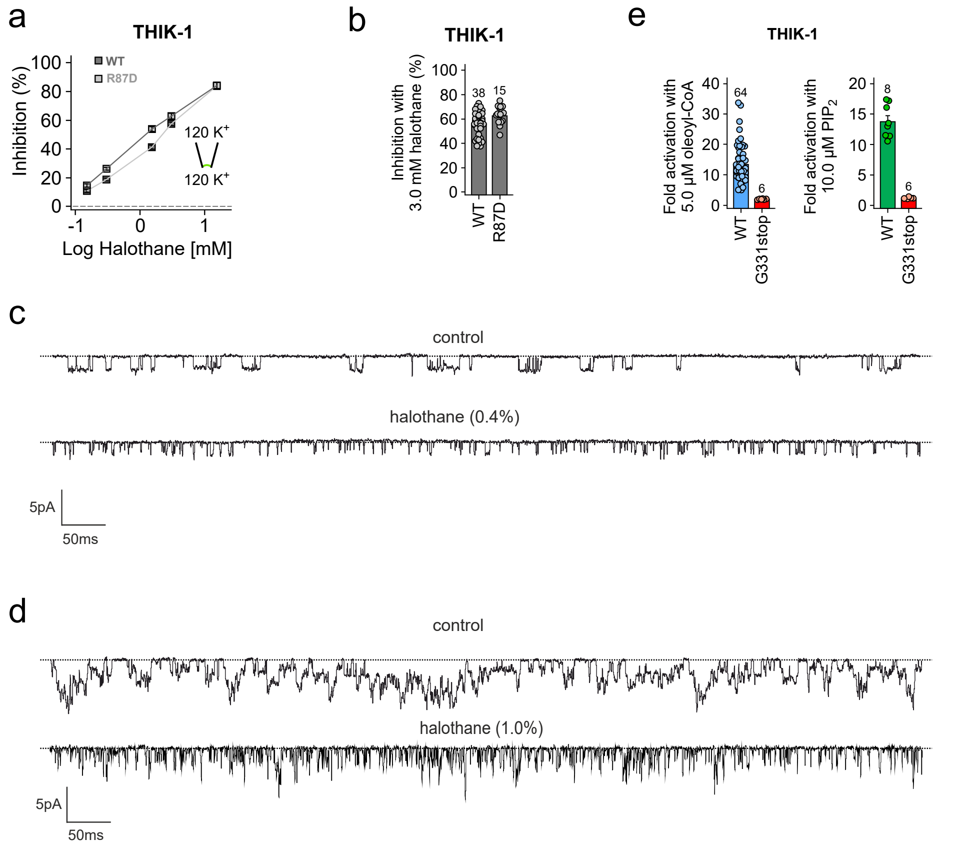

**Figure S5. Mechanisms of halothane inhibition and lipid activation of THIK-1**. **a,b,** Analysis of halothane inhibition for WT and THIK-1 R87D mutant channels, respectively showing that the R87D channel retains similar sensitivity to WT THIK-1. **c,d,** Excised patch recordings of 1 s duration at -100 mV of single WT THIK-1 channels in the absence (control) and presence of 0.4 % (c) or 1.0 % halothane (d). The dotted lines represent zero current levels. Halothane induces destabilisation (shortening) of both open and closed states. Thus, because destabilisation of channel closings increases $P_O$, it must be the destabilisation of channel openings that drives the inhibitory effect of halothane. Consistent with this, higher concentrations of halothane (panel d) induced a further, more dramatic reduction in the duration of openings, resulting in "flickery" single-channel kinetics and an even greater decrease in channel $P_O$ **e,** Analysis of fold activation of WT and G331x THIK-1 channels with 5.0 µM oleoyl-CoA and 10.0 µM $PIP_2$, respectively.
