## Supplementary material for "CryoEM Structure of the human THIK-1 K2P K^+^ Channel Reveals a Lower ‘Y-gate’ Regulated by Lipids and Anaesthetics": Higher Resolution Figures

### Figure 1

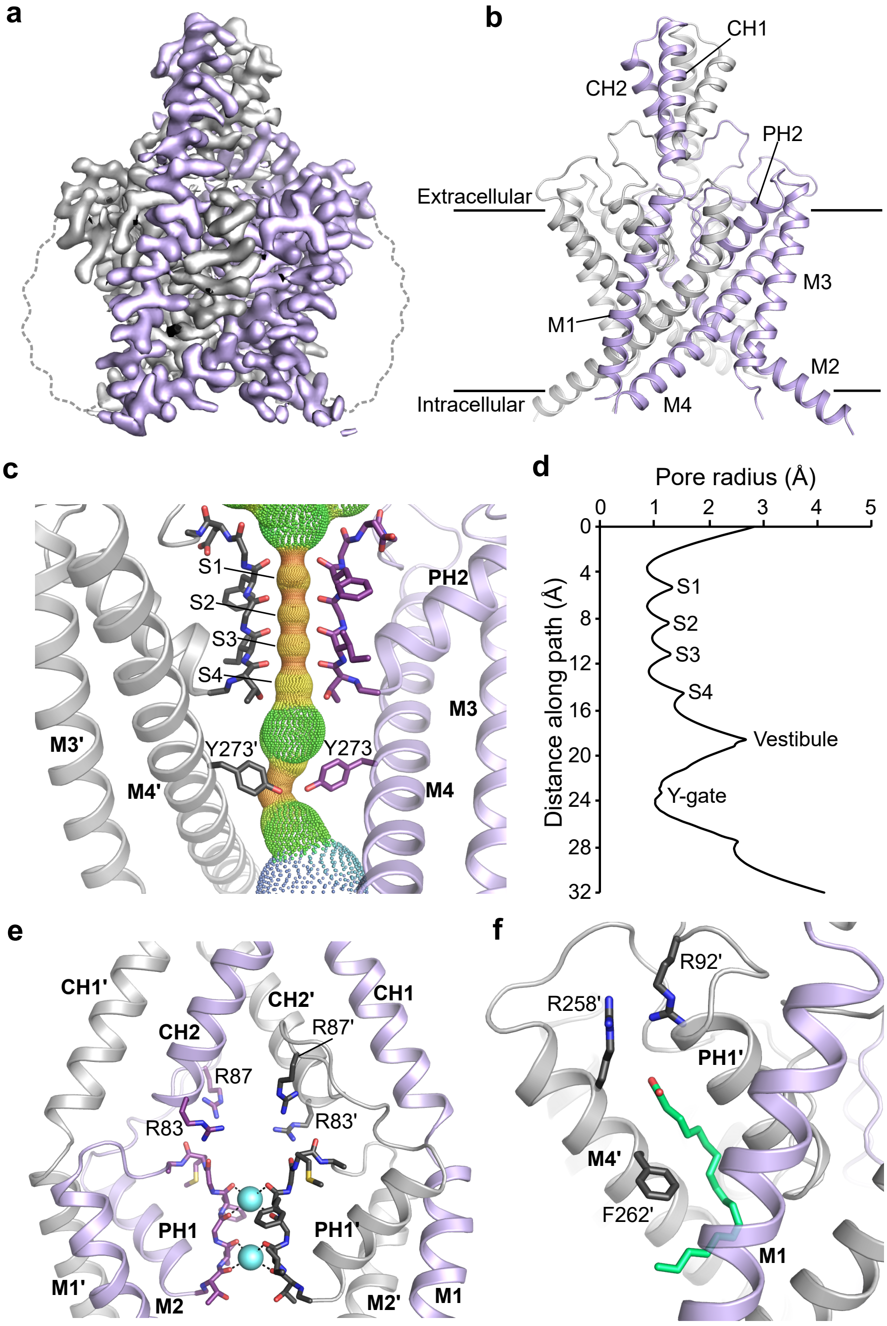


**Figure 1. Structure of THIK-1. a**, Sharpened cryoEM map viewed from the side with the density for THIK-1 channel subunits in gray and purple. The approximate position of the detergent micelle is outlined as a dotted gray line. **b**, Structure of THIK-1 coloured as in Panel a with the M1-M4, Pore and cap helices labelled. **c,** HOLE profile through the channel pore with the selectivity filter (S1-S4 sites) and the constriction formed by the lower tyrosine 273 (Y-gate) site highlighted. For clarity, M1, M2 and PH1 have been hidden. **d,** Pore radius of the THIK-1 channel interior as distance function along the ion permeation pathway. **e,** Ion exit pathway at the extracellular site of the selectivity filter is lined by a cluster of positively-charged residues (R83 and R87 from both subunits). M3, M4 and PH2 have been hidden for clarity. **f,** The K2P modulator pocket, showing a lipid bound at the inter-subunit interface of M4, PH1 and M1. Key residues in close proximity are highlighted as sticks.

### Figure 2

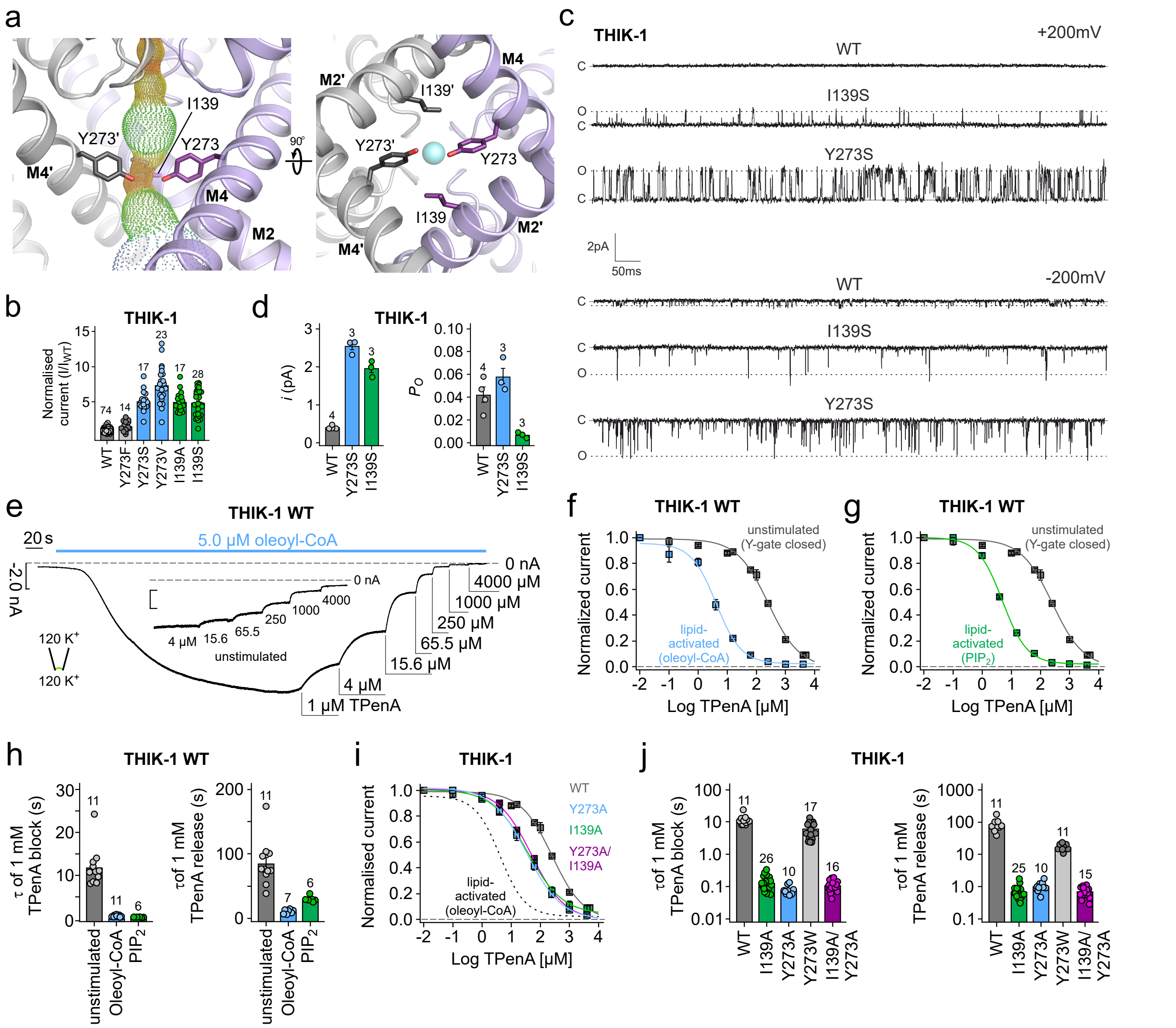


**Figure 2. THIK-1 Y-gate regulated by lipids. a,** The Y-gate viewed from the side (M2' hidden for clarity), and from the bottom, showing residue I139 in the same horizontal level as Y273 and also part of the constriction formed by the Y-gate. **b,** Relative whole-cell current amplitudes of WT THIK-1 and channels with mutations in the Y-gate. All currents are normalised to WT. **c,** Cell-attached recordings of 1s duration at ± 200mV containing single WT, I139S and Y273S THIK-1 channels, as indicated. The closed (c) and open channel (o) levels are indicated. **d,** Comparison of single-channel open probability (Po) and single channel current amplitude (*i*) of WT, I139S and Y273S THIK-1 channels in cell-attached patches at -200mV. Numbers above the bars denote the number of experiments. **e,** Representative recording at -80 mV from an inside out patch containing WT THIK-1 channels with symmetrical K+ concentrations (120 mM [K+]) at pH 7.4. Channel currents were activated with 5 µM oleoyl-CoA) applied to the intracellular side of the membrane and then inhibited dose-dependently with TPenA as indicated. Inlay shows equal inhibition with TPenA at the unstimulated basal state of the channels. **f,g,** Analysis of the apparent affinity for TPenA in the indicated states from recordings in panel e showing increased TPenA sensitivity after lipid activation. h, Analysis of TPenA kinetics for block and release of WT THIK-1 in unstimulated and lipid-activated states. **i-j,** Analysis of apparent TPenA affinity (i) and kinetics (j) for WT and indicated THIK-1 mutants from recordings as in the inlay in panel e. All values are given as mean ± s.e.m with number (n) of individual recordings indicated above the bars.

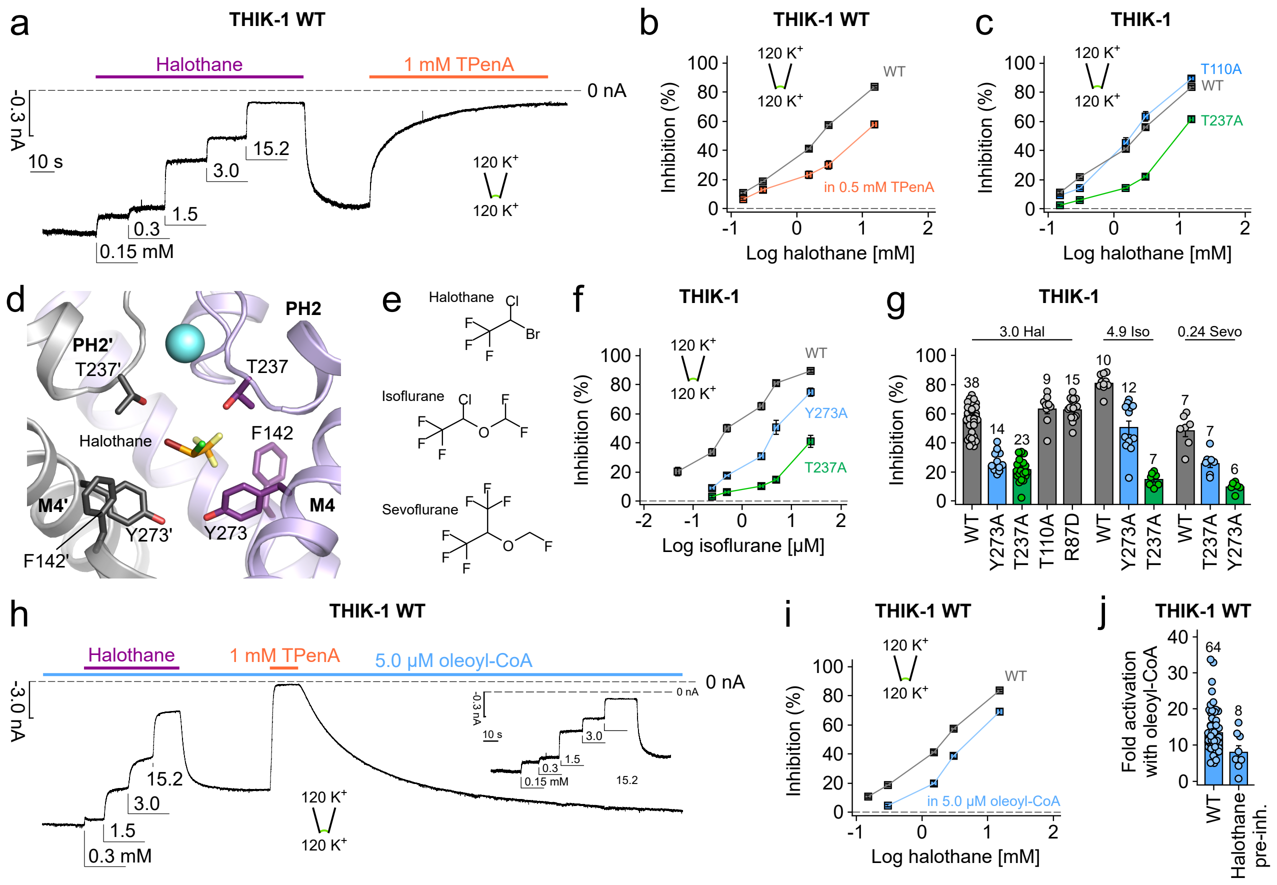


**Figure. 3 Inhibition by volatile anaesthetics involves both the filter and Y-gate. a,** Representative recording at -80 mV from an inside out patch containing WT THIK-1 channels with symmetrical $K^+$ concentrations (120 mM $[K^+]$ex./120 mM $[K^+]$int.) at pH 7.4. Channel currents were inhibited dose-dependently with increasing concentrations of halothane applied to the intracellular side of the membrane. Note, halothane effects can be washed and recovered and the currents inhibited with TPenA. **b,** Analysis of halothane inhibition for THIK-1 WT from recordings as in panel a in the absence (gray) and presence (orange) of 0.5 mM TPenA which produces ~ 80% block of initial currents. **c,** Analysis of halothane inhibition from recordings as in panel a for WT THIK-1 and indicated mutant channels. **d,** THIK-1 with docked halothane in the vestibule. Residues in close proximity are highlighted as sticks. For clarity, residues 121-138 in M2' as well as PH2' are not shown. **e,** Comparison of the structures of halothane, isoflurane and sevoflurane. **f,** Analysis of isoflurane inhibition of WT THIK-1 and channel mutants (*i.e.*, Y273A and T237A). **g,** Summary of volatile anaesthetic inhibition with either 3.0 mM halothane, 4.9 mM isoflurane and 0.24 mM sevoflurane for WT THIK-1 and mutant channels as indicated. **h,** Representative recording under conditions as in panel a showing dose-dependent halothane inhibition for THIK-1 channels activated with 5.0 μM oleoyl-CoA. **i,** Analysis of halothane inhibition in the absence and presence of 5.0 μM oleoyl-CoA from recordings as in panels a and h. **j,** Fold activation of WT THIK-1 with 5.0 μM oleoyl-CoA in the absence and presence of 15.2 mM halothane. Values are given as mean ± s.e.m with number (*n*) of individual recordings indicated above the bars.

Figure 4

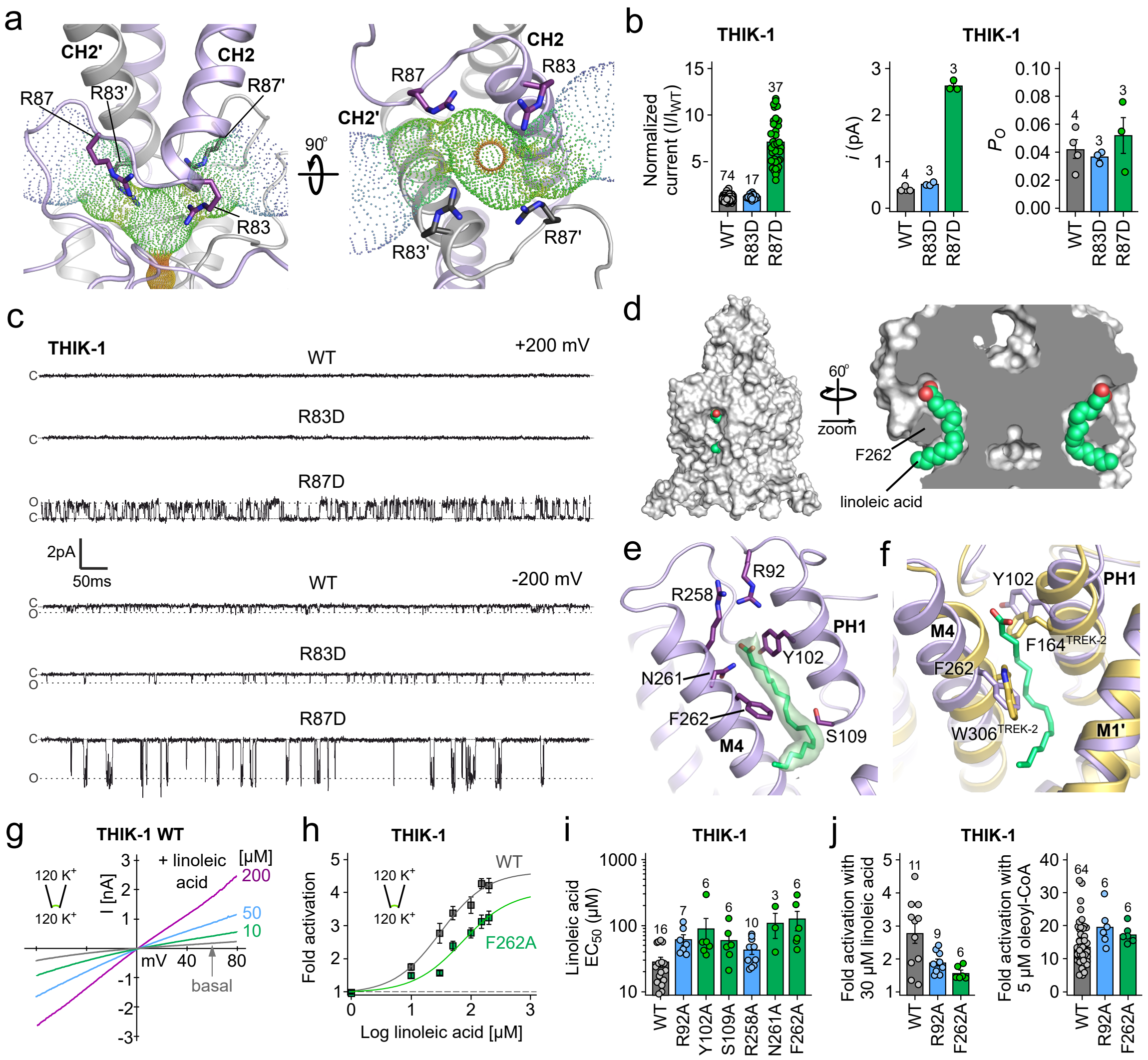


**Figure 4. Regulation of THIK-1 activity by charged residues and lipids. a,** Bifurcated extracellular ion exit pathway for $K^+$ showing the orientation of positively-charged residues in that region (R83 and R87). **b,** Left: relative whole cell current amplitudes of WT and mutant THIK-1 channels (R83D and R87D). Right: Analysis of the single channel amplitudes (*i*) and open probability ($P_O$) from $n \geq 3$ separate recordings as shown in panel c for WT and mutant THIK-1 channels in cell-attached patches at -200mV. **c,** Representative single channel recordings of WT THIK-1 compared to R83D and R87D mutant channels. Recordings shown at ± 200 mV in cell-attached configuration. The closed (c) and open (o) channel levels are shown. **d,** Surface representation of THIK-1 and cutaway showing two linoleic acid molecules (green) in the curved lipid binding pocket. **e,** The lipid binding pocket, with residues that have been mutated for functional studies highlighted as sticks. **f,** Structural overlay of THIK-1 (purple) with TREK-2 (yellow) showing position of F262 and Y102 in THIK-1 with equivalent W306 and F164 in TREK-2 which rotate during channel activation. **g,** Representative recording from an inside out patch containing WT THIK-1 with symmetrical $K^+$ concentrations (120 mM [$K^+$]) at pH 7.4. Channel currents were activated with indicated concentrations of linoleic acid. **h,** Analysis of fold activations from recordings as in panel g for WT and F262A mutant channels. **i,** $EC_{50}$ values from dose-response curves as in panel h for WT and mutant THIK-1. **j,** Fold activation of WT and R92A and F262A mutant THIK-1 channels, respectively with either 30μM linoleic acid or 5μM oleoyl-CoA. Values are given as mean ± s.e.m with number (*n*) of individual recordings indicated above the bars.
